# Suppression of de novo lipogenesis and dietary PUFA supplementation inhibit prostate cancer progression

**DOI:** 10.64898/2026.03.25.714193

**Authors:** Silvia D. Rodrigues, Caroline Fidalgo Ribeiro, Giuseppe N. Fanelli, Isadora Ferrari Teixeira, Alessandro Barberis, Hubert Pakula, Pier Vitale Nuzzo, Filippo Pederzoli, Fabio Socciarelli, Sara Bleve, Jeanne Jiang, Jonas Dehairs, Guilherme Henrique Tamarindo, Giorgia Zadra, Lisa M. Butler, Stephen R. Plymate, David W. Goodrich, Johannes V. Swinnen, David M. Nanus, Massimo Loda

**Affiliations:** Department of Pathology and Laboratory Medicine, Weill Cornell Medical College, New York, NY, USA; Department of Oncology, Pisa University Hospital, Division of Pathology, Department of Translational Research and New Technologies in Medicine and Surgery, University of Pisa, Pisa, Italy; Department of Medical Oncology, IRCCS Istituto Romagnolo per lo Studio dei Tumori (IRST) “Dino Amadori“, Meldola, IT; Laboratory of Lipid Metabolism and Cancer, Department of Oncology, Leuven Cancer Institute (LKI) and Leuven Institute for Single Cell Omics (LISCO), KU Leuven, Leuven, Belgium; Brazilian Biosciences National Laboratory, Brazilian Center for Research in Energy and Materials (CNPEM), Campinas, São Paulo, Brazil; Institute of Molecular Genetics, National Research Council (CNR-IGM), Pavia, Italy; South Australian Immunogenomics Cancer Institute, The University of Adelaide, Adelaide, SA, Australia; South Australian Health and Medical Research Institute, Adelaide, SA, Australia; Geriatric Research Education, and Clinical Center, Veterans Affairs Puget Sound Health Care System, Seattle, Washington, USA; Veterans Affairs Puget Sound Health Care System, Seattle, Washington, USA; Roswell Park Cancer Institute, Buffalo, NY, USA; Department of Medicine, Weill Cornell Medicine and the Meyer Cancer Center, New York, NY, USA; Department of Medical Oncology, Dana-Farber Cancer Institute, Harvard Medical School, Boston, MA; Nuffield Department of Surgical Sciences, Lincoln College, University of Oxford, Oxford, UK

**Keywords:** prostate cancer, lipid metabolism, diet, fatty acids, cancer metabolism, lipogenesis

## Abstract

Prostate cancer progression is characterized by dysregulated lipid metabolism, with activation of fatty acid synthase (FASN), the rate-limiting step in *de novo* lipogenesis (DNL), resulting in significant accumulation of saturated lipids. Here, we show that pharmacologic FASN inhibition creates a metabolic state that increases reliance on exogenous polyunsaturated fatty acids (PUFAs). Inhibition of FASN profoundly alters membrane phospholipid composition, driving compensatory incorporation of PUFAs into membrane phospholipids, thus increasing susceptibility to lipid peroxidation and oxidative damage. Combined FASN inhibition and PUFA exposure induce mitochondrial hyperpolarization and enhance lipid peroxidation in both hormone-sensitive and castration-resistant prostate cancer models, resulting in increased reactive oxygen species production, ferroptosis, as well as apoptosis. Marked inhibition of growth in castration-resistant human and murine prostate cancer organoids is achieved *ex vivo*. In genetically engineered, DNL-reliant Hi-Myc mice, a diet enriched in PUFAs significantly inhibited invasive carcinoma compared to a saturated fat-enriched diet. Thus, environmental PUFAs modulate and enhance the therapeutic efficacy of FASN-targeted strategies. These findings set the stage for pharmacologic and dietary intervention in prostate cancer patients.

## 2. Introduction

Metabolic reprogramming is a hallmark of cancer^1^. In prostate cancer, neoplastic cells exhibit distinct metabolic alterations compared to normal prostate epithelium, most notably a strong dependence on fatty acid (FA) metabolism^2^. As a result, upregulation of lipogenesis plays a pivotal role in both tumor initiation and progression ^3–5^.

Fatty acid synthase (FASN), the rate-limiting enzyme of DNL, is a central driver of this metabolic shift. FASN overexpression occurs early in tumorigenesis and persists throughout disease progression, reaching the highest expression levels in bone metastases^6–8^. FASN catalyzes the *de novo* synthesis of palmitate (16:0), a long-chain saturated fatty acid, from acetyl-CoA and malonyl-CoA^9^. Palmitate serves as a precursor for the generation of a spectrum of saturated fatty acids (SFAs) and monounsaturated fatty acids (MUFAs) through elongation and desaturation pathways^3,10^. These FAs are indispensable for membrane biosynthesis, energy metabolism and modulation of oncogenic signaling via post-translational modifications^11–13^. Pharmacologic or genetic inhibition of FASN suppresses tumor growth, induces cell death, enhances the efficacy of androgen receptor-targeted therapies, and synergizes with poly ADP-ribose polymerase (PARP) inhibitors by promoting DNA damage; all these effects can be rescued by exogenous palmitate supplementation^4,7,14^. An obesogenic, high-fat diet amplifies MYC-driven transcriptional programs, promoting histone hypomethylation, glycolytic flux, and lactate accumulation^15^. Notably, switching from a high- to low-saturated-fat diet attenuates MYC signaling in murine models^15^. Consistent with a role for dietary fat quality in prostate cancer outcomes, greater adherence to healthy dietary patterns after diagnosis, particularly those enriched in unsaturated fats and reduced saturated fats, was associated with improved overall survival, while a higher saturated fat intake has been linked to worse outcomes in this population^16^. Interestingly, inhibition of FA uptake (e.g., via CD36, also known as Fatty Acid Translocase (FAT), which facilitates the rapid uptake of FAs into cells) reduces proliferation and aggressiveness in preclinical prostate cancer models, underscoring the reliance on external lipid sources when intrinsic biosynthesis is disrupted^17^.

Unlike SFAs and MUFAs, polyunsaturated fatty acids (PUFAs, ≥ 2 acyl chain double bonds) cannot be fully synthesized *de novo* in mammalian cells. The parent compounds must be obtained exogenously, primarily through dietary intake. When endogenous lipid biosynthesis is impaired, the composition of the extracellular fatty acid pool becomes an important determinant of cell fate. Whereas exogenous SFAs and MUFAs limit lipid peroxidation and ferroptosis, PUFA enrichment increases oxidation-prone phospholipid species and enhances ferroptotic vulnerability^18^. This adaptive response indicates a critical metabolic shift, highlighting the reliance of cells on external lipid sources when internal lipid biosynthetic pathways are disrupted. Due to their chemical structure, PUFAs are highly susceptible to attack by reactive oxygen species (ROS). This vulnerability leads to the formation of lipid peroxyl radicals (lipid-ROS), which initiate and propagate a chain reaction in adjacent membrane lipids^19,20^. This phenomenon amplifies oxidative damage within the cell and can ultimately compromise membrane integrity, triggering cell death pathways, including ferroptosis, particularly in cancer cells under metabolic stress^19,21–24^.

Taken together, these findings suggest that simultaneous inhibition of cancer cell– autonomous SFA/MUFA synthesis and enrichment of the tumor lipid milieu with PUFAs may induce a tumor-specific lipid remodeling program with consequences for cell growth and death. Here, we investigated both *in vitro* and *in vivo* whether exogenous PUFAs, in combination with pharmacological inhibition of FASN could be exploited to affect prostate cancer survival.

## 3. Material and Methods

### Reagents

The following reagents were used in the study: IPI-9119 (Infinity Pharmaceuticals) and TVB-2640 (*Sagimet* Biosciences) for *in vitro* treatment, TVB-3664 (*Sagimet* Biosciences) for *in vivo* treatment, Ferrostatin-1 (#S7243, Selleck Chemicals), palmitate-BSA FAO substrate (#102720-100, Agilent Technologies), Matrigel matrix phenol red-free growth factor reduced (Corning, #ref 356231), PUFAs: *cis*-4,7,10,13,16,19-Docosahexaenoic acid (DHA, #D2534, Sigma-Aldrich, USA), *cis*-7,10,13,16,19-Docosapentaenoic acid (DPA, #D1797, Sigma-Aldrich), *cis*-5,8,11,14,17-Eicosapentaenoic acid (EPA, #E2011, Sigma-Aldrich).

### FASN inhibitors

FASN activity was pharmacologically inhibited using selective small-molecule FASN inhibitors (FASNi). Distinct inhibitors were selected according to experimental context and formulation suitability. For in vitro studies in prostate cancer cell lines, IPI-9119 (100 nM) was used. MSK-PCa3 organoids were treated with IPI-9119 at 500 nM. For murine prostate organoid experiments, TVB-2640 (clinically formulated, GMP-grade) was used at 500 nM. In vivo studies were performed using TVB-3664 (10 mg/kg, 5 days per week), a close structural analog of TVB-2640 optimized for systemic exposure and tolerability in mice. TVB-3664 was dissolved in PEG400 to prepare a 20 mg/mL stock solution. Stocks were prepared weekly and stored protected from light at room temperature. Immediately prior to administration, the stock was diluted in sterile water to obtain a final formulation of 30% PEG400 and 70% water (v/v). Mice received TVB-3664 by oral gavage at 10 mg/kg, with dosing volume adjusted according to body weight. Vehicle-treated controls received the corresponding PEG400:water formulation without compound. IPI-9119 inhibits the β-ketoacyl reductase domain of FASN^4,14^, whereas TVB-2640 and TVB-3664 target the thioesterase domain^25^. All inhibitors suppress de novo lipogenesis. Importantly, key phenotypes observed in vitro were recapitulated using structurally distinct FASN inhibitors, supporting the on-target mechanism and enhancing translational confidence.

### Cell culture and treatments

Human prostate cancer cell lines, including hormone-sensitive LNCaP (prostate adenocarcinoma, lymph node metastasis, RRID:CVCL_1379), and castration-resistant 22Rv1 (prostate adenocarcinoma, RRID:CVCL_1045), and C4-2 (androgen-independent LNCaP subline, RRID: CVCL_4782), were obtained from the American Type Culture Collection (ATCC). Cells were maintained in RPMI 1640 medium (Gibco, Cat# 11835-030) supplemented with 10% fetal bovine serum (FBS, Sigma, Cat#F4135) and incubated at 37°C in a humidified atmosphere containing 5% CO2. All cell lines were authenticated by ATCC through short tandem repeat (STR) profiling and confirmed to be mycoplasma-free. To ensure consistency, cells were used only up to passage 50. DHA was used at 100 μM alone or in combination with FASN inhibitors. EPA (120 μM), DPA (100 μM) were used alone or in combination with FASNi. Ferrostatin-1 was used at 10 μM in complete medium.

### Measurement of cellular bioenergetics and mitochondrial function

Following 6 days of incubation with FASNi, cells were collected with trypsin–EDTA 0.25%, counted, and reseeded at a density of 2 × 10^4^ cells in Seahorse XF^96^ V3 PS Cell Culture Microplates (#101085–004, Agilent Technologies Inc, USA) using the same treatment medium to maintain consistent FASNi concentrations. Mitochondrial function was assessed using the Seahorse XF Cell Mito Stress Test assay (#103015–100, Agilent Technologies Inc, USA) according to the manufactures’ instructions. Briefly, Oligomycin (1 µM), FCCP (2 µM) and Rotenone/Antimycin A (0.5 µM) were loaded into the designated cartridge ports. FCCP concentration and cell density were optimized through prior titration experiments. Before the assay, culture medium was replaced with warm and freshly prepared Seahorse XF Base Medium (#103681–100, Agilent Technologies Inc, USA), supplemented with 1 mM pyruvate, 2 mM glutamine and 10 mM glucose (pH 7.4). Plates were incubated at 37°C in a non-CO₂ incubator for 45 min prior to measurement. Oxygen consumption rate (OCR) and extracellular acidification rate (ECAR) were determined using the Seahorse XFe96 Analyzer (Agilent Technologies Inc., USA). At the end of the assay, cells were lysed with RIPA buffer, and total protein content was quantified by BCA assay for normalization. Data analysis was performed using Wave software (v2.6.3, Agilent). Parameters were calculated as follows: Basal Mitochondrial Respiration = (Late rate measurement before first injection) – (Non-mitochondrial respiration rate); Maximal Mitochondrial Respiration = (Maximum rate measurement after FCCP injection) – (Non-mitochondrial respiration rate); Proton Leak = (Minimum rate measurement after Oligomycin injection) – (Non-mitochondrial respiration); ATP-linked Production = (Late rate measurement before Oligomycin injection) – (Minimum rate measurement after Oligomycin injection); Spare Capacity = (Maximal Mitochondrial Respiration)—(Basal Mitochondrial Respiration); Non-mitochondrial respiration rate = Minimum rate measurement after Rotenone/ Antimycin A injection. ECAR was recorded concurrently during the assay. Three independent experiments were performed with at least three technical replicates per condition (n = 3). Data are presented as mean ± SEM, expressed as pmol O₂/min/protein for OCR and mpH/min/protein for ECAR.

### [¹⁴C]-DHA uptake assay

Cells were seeded at a density of 1.5 × 10⁵ cells per 35-mm dish and treated with DMSO or the FASN inhibitor IPI-9119 (100 nM) for 6 days. For uptake measurements, cells were incubated with 2 µL of [¹⁴C]-DHA (0.5 µCi/µL; ARC0380) for 3 h at 37 °C. Following incubation, cells were washed with PBS, detached with trypsin, and pelleted by centrifugation (300 × g, 5 min, RT). Pellets were washed once in PBS and lysed in 200 µL RIPA buffer on ice for 15 min. Lysates were cleared by centrifugation (14,000 × g, 10 min, 4 °C), and 150 µL of the supernatant was mixed with 10 mL of scintillation fluid (PerkinElmer) for β-emission counting using a Tri-Carb 2910TR Liquid Scintillation Counter. Radioactivity was expressed as counts per minute (cpm) and normalized to total protein content. Data represent three independently performed experiments.

### Competitive radiolabeled fatty-acid uptake assay

Cells were seeded at a density of 1.5 × 10⁵ cells per well in 6-well plates and treated with DMSO or the FASN inhibitor IPI-9119 (100 nM) for 6 days. Cells were then incubated for 3 h with one of the two: (1-PUFA labeled) 0.45 µM [14C]-DHA (ARC-0380; 0.1 µCi/µL) plus 0.45 µM unlabeled palmitate or (2-SFA labeled) 0.45 µM unlabeled DHA (Sigma D2534) plus 0.45 µM [14C]-palmitate ([1-14C]-palmitic acid, ARC-01762A; 0.5 µCi/µL). Radiolabeled FAs were conjugated to 10% fatty acid–free BSA at 37 °C prior to use. After incubation, cells were washed with PBS, lysed in RIPA buffer, and 150 µL aliquots were analyzed by liquid scintillation counting. Radioactivity (cpm) was normalized to total protein content from parallel non-radioactive wells. Data represent three independently performed experiments.

### Measurement of ROS

Intracellular mitochondrial superoxide anion and mitochondrial membrane potential levels were determined using MitoSOX^TM^ Red (#M36008, Life Technologies) and JC-1 dye (#ab113850, Abcam), respectively, according to the manufacturer’s instructions. Briefly, cells were seeded at 1.5×10^5^ into a 6-well plate. After 40 hours, the medium was replaced, and DMSO, FASN inhibitors, and DHA were incubated in fresh media for 6 days. Following the treatment, cells were collected and counted, then incubated with dyes (5 µM Mitosox; 10 µM JC-1) for 30 min at 37°C. Fluorescence was measured at 510/580 nm wavelengths for MitoSox, and 535/590 nm and 475/530 nm to quantify aggregated and monomeric forms in JC-1, respectively. All values were normalized to the total number of viable cells.

### Lipid peroxidation

After 6 days of treatment with FASN inhibitors and PUFAs, cells were collected, washed with fresh medium, and incubated with 5 μM BODIPY 581/591 C11 (#D3861, Thermo Fisher Scientific), at 37°C for 30 min. Cells were washed with PBS, resuspended in fresh phenol-free medium, and transferred to a 96-well plate. Fluorescence emission shift was measured by reading the probe’s excitation/emission at 571/591 nm (oxidized) and 488/510 nm (reduced). The oxidized to reduced probe ratio was normalized to the total number of viable cells.

### Protein carbonylation

Protein carbonylation levels were assessed using total protein lysates prepared in RIPA buffer. Carbonyl groups were derivatized with 2,4-dinitrophenylhydrazine (DNP) and detected by ELISA, following a protocol adapted from Buss et al.^26^. Detection was performed using an anti-DNP antibody (Molecular Probes Cat# A-6430, RRID: AB_221552; dilution 1:2700) incubated for 1 hour at 37°C, followed by HRP-conjugated secondary antibody treatment. Untreated cells exposed to H2O2 (100 μM,1 hour) were used as a positive control for protein oxidation.

### Immunoblotting

Cell lysates were prepared in RIPA buffer supplemented with protease (PhosSTOP Roche Cat#04906837001) and phosphatase inhibitors (cOmplete Mini, EDTA-free Roche Cat#11836170001). Protein concentration was determined using the Bradford Protein Assay (Bio-Rad). Equal amounts of total lysate (15 µg) were resolved on precast Tris-Glycine gels (Invitrogen) and transferred to nitrocellulose membranes. The following antibodies were used: BiP (CST Cat# 3177, RRID: AB_2119845), ACSL4/FACL4 (Abcam Cat# ab155282, RRID: AB_2714020), GPX4 (CST Cat# 52455, RRID: AB_2924984), AR (Abcam Cat# ab108341, RRID:AB_10865716), and Vinculin (Sigma-Aldrich Cat# V9131, RRID:AB_477629). HRP-conjugated secondary antibodies included Goat Anti-Rabbit IgG (Bio-Rad Cat# 170-6515, RRID: AB_11125142) and Goat Anti-Mouse IgG (Bio-Rad Cat# 170-6516, RRID:AB_2921252). Blots were developed with Pierce ECL and imaged on a G-Box Chemi XRQ system (Syngene). Band intensities were quantified using GeneTools software (Syngene).

### Lipidomics

Lipidomic profiling was performed at the KU Leuven lipidomics core facility Lipometrix (KU Leuven, Belgium) using a liquid chromatography-electrospray ionization tandem mass spectrometry (LC-ESI/MS/MS) platform. Samples were analyzed on a Nexera X2 UHPLC system (Shimadzu) coupled with an AB SCIEX 6500+ QTRAP, a hybrid triple quadrupole/linear ion trap mass spectrometer, as previously described^27^.

### Genetically engineered mouse models and dietary interventions

FVB Hi-Myc male mice (MGI:5486199), which overexpress the human c-MYC transgene in the prostatic epithelium (ARR2PB-Myc-PAI), were obtained from the National Cancer Institute Mouse Repository at the Frederick National Laboratory for Cancer Research. Experimental breeders were selected, and cohorts were established (n=10 per time point) after randomization and assigned to different dietary regimens differing in fat composition (Supplementary Table 1). Upon weaning (21 days of age), male mice were fed on either a saturated/monounsaturated fatty acid HSD diet (HSD; Tekland/Envigo: TD.230313) or a polyunsaturated fatty acid diet (HPD; Tekland/Envigo: TD.230355) and maintained on these diets for 12 or 20 weeks. Diets were purified, isocaloric (4.1 kcal/g), macronutrient-matched (15% protein, 55% carbohydrate, 30% fat kcal), differing only in FA composition. The FASN inhibitor TVB-3664 (10 mg/kg 5 days per week) or vehicle control was administered starting at week 7 (12-week cohort) or at week 14 (20-week cohort), for 5 and 6 weeks, respectively. Food was replenished weekly, and mice were weighed every three weeks before euthanasia. Mice were housed under a 12-hour light/dark cycle with unrestricted access to food and water at Weill Cornell Medicine Animal Resources. The Weill Cornell Medicine Institutional Animal Care and Use Committee (IACUC) reviewed and approved the protocol (protocol number: 2023-0023) in accordance with the Animal Welfare Act, adhering to the ethical guidelines of the RARC at Weill Cornell Medicine. Tumor burden in male Hi-Myc mice did not result in any observable adverse effects before the experimental endpoint (i.e., 20 weeks of age).

### Histopathology

At the endpoints, mice were weighed and euthanized. Prostates were collected and weighed. Anterior (AP), dorsolateral (DLP), and ventral prostate (VP) lobes were dissected as previously described^28^, and separated into right and left hemi-prostates, individually weighed, measured. The right hemi-prostate was fixed in 10% neutral-buffered formalin for paraffin embedding, whereas the left hemi-prostate was snap-frozen on dry ice and stored at −80°C for subsequent lipidomic analyses. Formalin-fixed paraffin-embedded (FFPE) murine blocks were sectioned at 5 micrometers and mounted on Leica-charged slides. Consecutive sections were stained for hematoxylin and eosin (H&E). All stained slides were imaged using an Akoya Biosciences PhenoImager HT 2.1.0. For each treatment condition, the H&E-stained prostate section was digitized at 20x magnification using the Leica Aperio GT450 slide scanner (Leica Biosystems, Wetzlar, Germany). All histopathologic evaluations were carried out by a board-certified pathologist (GNF). For each timepoint, comparisons between corresponding lobes across dietary groups were performed using the non-parametric Mann–Whitney U test. Statistical analyses were conducted in GraphPad Prism (version 10.4.1; GraphPad Software, San Diego, CA). Histopathological processing was conducted in collaboration with the Weill Cornell Medicine Center for Translational Pathology (RRID:SCR_027311) and the Multiparametric In Situ Imaging Laboratory (RRID:SCR_024591) Core Facilities.

### Organoid establishment and culture

Hi-Myc prostate cancer organoids were established from the prostates of 6-month-old Hi-Myc mice. Prostate tissues were digested in Advanced DMEM/F12 (Gibco, Cat#12634028) containing 1.5 mg/mL Collagenase II, 1000 U/mL Hyaluronidase VIII (Thermo Fisher Scientific), and 10 μM Y-27632 (Dihydrochloride, STEMCELL Technologies, Cat#72304) for 1 hour at 37°C with continuous agitation on an orbital shaker (Max Q 8000, Thermo Scientific at 250 rpm). After centrifugation at 150 g (Beckman Coulter, Allegra X-30R) for 5 min at 4°C, cells were resuspended in TrypLE (Gibco, Cat#12604-013) supplemented with 10 μM Y-27632, incubated for 15 min at 37°C, and neutralized in DMEM/F12 with 0.05% FBS. Single-cell suspensions were obtained by filtering through 70 μm and 40 μm strainers (BD Biosciences). Cells were washed once in mouse organoid medium^29,30^ and resuspended in growth factor-reduced Matrigel (Corning, 354230) diluted 1:1 with organoid medium. Approximately 1 mL of the Matrigel-cell mix was used per mouse. Droplets were plated onto 10 cm dishes, solidified for 15 min at 37°C, and overlaid with 10 mL of prewarmed organoid medium. Cultures were maintained at 37°C in 5% CO₂, and the medium was replaced every 3 days. Organoids were passaged weekly at a 1:3 ratio by dissociation in cold organoid medium, centrifugation at 750 g for 5 min at 4°C, and digestion in TrypLE Express for 15 min at 37°C. Cells were washed and replated in fresh Matrigel/media mix. Organoids were cryopreserved in 90% FBS and 10% DMSO and stored at −150°C.

The human MSK-PCa3 prostate cancer organoid line was kindly provided by Dr. Yu Chen (Memorial Sloan-Kettering Cancer Center, NYC). MSK-PCa3 and Hi-Myc organoids were dissociated into single cells using TrypLE for 20 min at 37°C. For organoid formation, 5 × 10³ cells were resuspended in 20 µL of growth factor-reduced Matrigel and appropriate human organoid medium^31^ (1:1, v/v) and seeded in 12-well plates. Organoids were treated with FASN inhibitors, PUFAs, or DMSO as a control, as previously described^4,14,32^. Medium was replaced every three days over 15 days. Organoid images were captured using a THUNDER Imager LEICA DMi8 Microscope (LEICA, Germany) with LAS X software (version 3.7.4.23463). Organoid number and diameter were quantified using ImageJ software (National Institutes of Health), with measurements conducted manually.

### Statistical analysis

For statistical analyses, GraphPad Prism 10 was used. Multiple comparisons were performed using one-way ANOVA with Tukey’s multiple-comparison test, and adjusted P values were reported. Multiple independent comparisons were performed using two-way ANOVA with Šídák’s or Dunnett’s multiple-comparison test, and adjusted P values were reported in the figure legend. All the experiments were performed using at least three independent biological replicates. For clarity, statistical annotations displayed in the main figure 5 indicate comparisons between the combination treatment and single-agent conditions.

For lipidomics analysis, peak integration was done using MultiQuantTM 3.0.3 software. Lipid species signals were corrected for isotopic contributions with Python Molmass 2019.1.1, and quantified based on internal standard signals, adhering to the guidelines of the Lipidomics Standards Initiative (LSI) (level 2 type quantification as defined by the LSI). Counts were then normalized to DNA content. A heat map of lipid species was generated using Morpheus (https://software.broadinstitute.org/morpheus/) from normalized values of the indicated lipid class, summed by acyl chain fatty acid. For lipidomic analyses of the *in vivo* datasets, involving simultaneous testing of multiple lipid species, false discovery rate (FDR) correction was applied to account for multiple hypothesis testing using the two-stage step-up procedure of Benjamini, Krieger, and Yekutieli as implemented in GraphPad Prism. Adjusted q values were used to determine statistical significance.

## 4. Results

### 4.1 Pharmacologic inhibition of FASN alters energy metabolism in hormone-sensitive and castration-resistant PCa cells

We previously demonstrated that FASN inhibition disrupts lipid biosynthesis, primarily by reducing the intracellular palmitate pool^4,14^. Given the central role of lipid metabolism in energy production, we first determined how FASN blockade affects the cellular bioenergetic landscape of hormone-sensitive (HS) and castration-resistant (CR) prostate cancer cells. We evaluated mitochondrial and glycolytic functions in HS (LNCaP; Figure 1A) and CR (C4-2; Figure 1B; and 22Rv1; Figure 1C) prostate cancer cell lines, which were treated with the FASN inhibitor (FASNi) for 6 days. FASN inhibition significantly impaired ATP-linked respiration and basal respiration in CR cells (left and middle panels, respectively). ECAR did not show a compensatory increase, indicating that FASN inhibition primarily reduces mitochondrial oxidative metabolism without a marked glycolytic shift.

**Figure 1.**
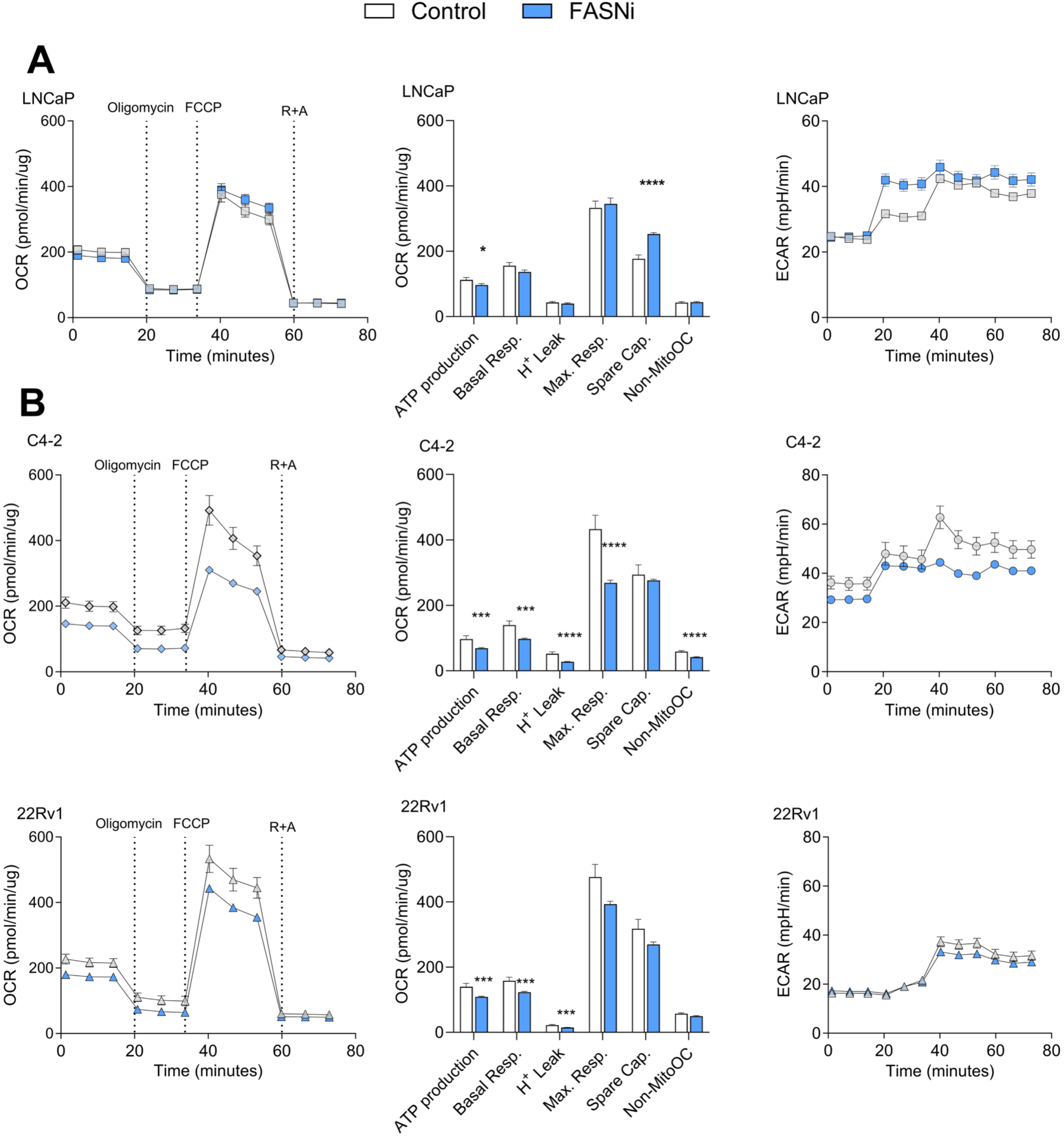
Pharmacologic FASN inhibition impairs mitochondrial oxidative metabolism in prostate cancer cells. (A-C) Oxygen consumption rate (OCR) and extracellular acidification rate (ECAR) in LNCaP, C4-2, and 22Rv1 cells treated with the FASN inhibitor IPI-9119 (100 nM) for 6 days, assessed using the Seahorse XF Cell Mito Stress Test. Quantification of ATP-linked respiration, basal respiration, proton leak, maximal respiration, spare respiratory capacity, and non-mitochondrial respiration was derived from OCR measurements. Data were normalized to total protein content per well and represent mean ± SEM from three independent biological replicates (n = 3). Statistical comparisons were performed using two-tailed unpaired t-tests with Welch’s correction. *P < 0.05; ***P < 0.001; ****P < 0.0001.

Taken together, these results, along with the known inhibition of β-oxidation secondary to malonyl-carnitine accumulation resulting from FASN inhibition^4^, show that FASN inhibition negatively impacts cellular energy metabolism, particularly in CR cells.

### 4.2 FASN blockade induces PUFA-enriched membrane remodeling and ferroptotic priming

FASN inhibition restricts endogenous production of SFAs and MUFAs, which are major components of membrane phospholipids^33^. In cells cultured in standard serum-containing medium without exogenous PUFA supplementation, lipid profiling revealed that pharmacological blockade of FASN markedly enriched PUFAs in major membrane phospholipid classes (Extended Data S1). Specifically, PUFA containing phosphatidylcholine (PC) and phosphatidylethanolamine (PE) species were increased across all prostate cancer cell lines analyzed (Figure 2A), consistent with increased reliance on exogenous lipid uptake when *de novo* fatty acid synthesis is suppressed. FASN inhibition also increased the ratio of PUFA-containing PC to PE species (Figure 2B). Changes in the PC ratio can affect membrane organization and ER homeostasis^34^, and may contribute to the endoplasmic reticulum (ER) stress associated with inhibition of DNL. Consistent with this, we previously have shown that inhibition of lipid synthesis induces ER stress, which can be rescued by palmitate supplementation^4,35^.

**Figure 2.**
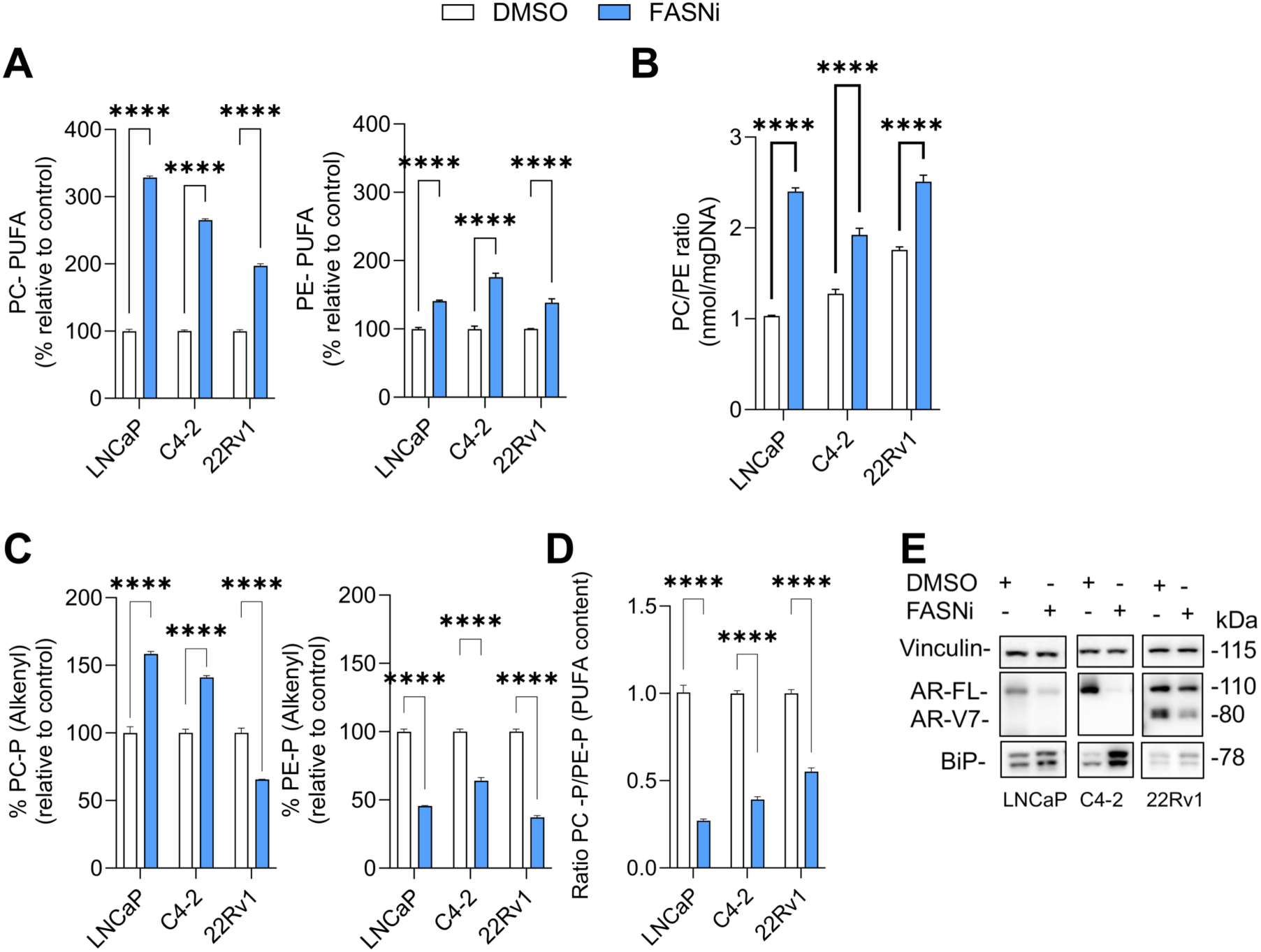
FASN inhibition induces PUFA-enriched membrane remodeling and alters lipid-associated stress signaling. (A–E)) Lipidomic analysis of phospholipid remodeling in LNCaP, C4-2, and 22Rv1 cells treated with FASNi (IPI-9119 100 nM) for 6 days under standard serum-containing conditions. (A) PUFA-containing phosphatidylcholine (PC-PUFA) species and PUFA-containing phosphatidylethanolamine (PE-PUFA) species were expressed relative to control. (B) Ether-linked (Alkenyls) PC-P and PE-P abundance expressed relative to control. (C) Ratio of total ether-linked PC to PE species (PC-P_total/PE-P_total). (D) Ratio of PUFA-containing ether-linked PC to PE species (PC-P_PUFA/PE-P_PUFA). (E) Immunoblot analysis of AR and BiP following FASN inhibition. Lipid species were normalized to DNA content and expressed relative to DMSO-treated control.

Inhibition of DNL also induced selective remodeling within the total ether-linked phospholipid pool independently of acyl-chain saturation or length. Total alkenyl-PC (PC-P_total) increased, whereas total alkenyl-PE (PE-P_total) decreased, resulting in a ratio imbalance (Figure 2C). Because PC- and PE-type ether phospholipids differ in their effects on membrane packing and curvature, this shift may alter membrane biophysical properties and contribute to lipid bilayer stress ^36,37^. In contrast, analysis restricted to PUFA-containing ether phospholipids showed a relative enrichment of PUFAs in PE-P, resulting in a lower PC-P_PUFA/PE-P_PUFA ratio (Figure 2D). PE-P species are preferentially enriched in long-chain PUFAs^38^, suggesting that this redistribution may increase the availability of oxidizable phospholipid substrate. Together, these data show that FASN inhibition induces distinct changes in membrane phospholipid composition, characterized by altered ether-linked phospholipid substrates. These changes were accompanied by induction of the ER stress marker BiP and modulation of AR expression following FASN inhibition (Figure 2E).

The enrichment of long-chain PUFA-containing phospholipids and redistribution within ether-linked species predict increased susceptibility to lipid peroxidation, a key initiating event in ferroptotic cell death. Because ferroptosis is driven by the iron-dependent accumulation of phospholipid hydroperoxides, we next examined ACSL4 and GPX4, central regulators of PUFA incorporation into membrane phospholipids and lipid peroxide detoxification, respectively. FASN inhibition increased ACSL4 expression, consistent with enhanced esterification of long-chain PUFAs into membrane phospholipids under DNL blockade. GPX4 protein expression was reduced, consistent with decreased antioxidant capacity to buffer phospholipid hydroperoxides (Figure 3A). To determine whether this lipid remodeling phenotype was followed by a broader, regulated cell death response, we next analyzed a curated panel of genes linked to ferroptosis, apoptosis, and necroptosis using a joint additive model adjusted for baseline cell-line differences (GSE114016, LNCaP and 22Rv1; and GSE224323, C4-2)^4,14^. This analysis revealed a ferroptosis-priming signature after FASN inhibition, with increased ACSL4 and LPCAT3, supporting enhanced activation and incorporation of oxidizable PUFA acyl chains into membrane phospholipids, together with increased transferrin receptor 1 (TFRC1 or CD71), consistent with elevated iron-import capacity (Figure 3B). Furthermore, FASN inhibition modulated transcripts associated with extrinsic apoptosis and necroptosis, including TNFRSF1A, CFLAR, XIAP, BID, BAX, BAK1, and MLKL, indicating broader remodeling of stress-death signaling. However, because transcript abundance does not establish pathway execution, we functionally assessed the contribution of ferroptosis versus apoptosis involvement. We performed rescue experiments using Ferrostatin-1 (Fer-1), a selective inhibitor of lipid peroxidation, or z-VAD-fmk (z-VAD), a pan-caspase inhibitor, in combination with FASN inhibition. Both Fer-1 and z-VAD partially restored cell viability (Figure 3C), indicating that FASN blockade engages overlapping lipid peroxide-dependent and caspase-dependent death mechanisms. Together with the lipidomic enrichment of PUFA-containing phospholipids and the Fer-1 rescue data, this transcriptional profile supports a model in which FASN blockade lowers the threshold for lipid peroxide–mediated death.

**Figure 3.**
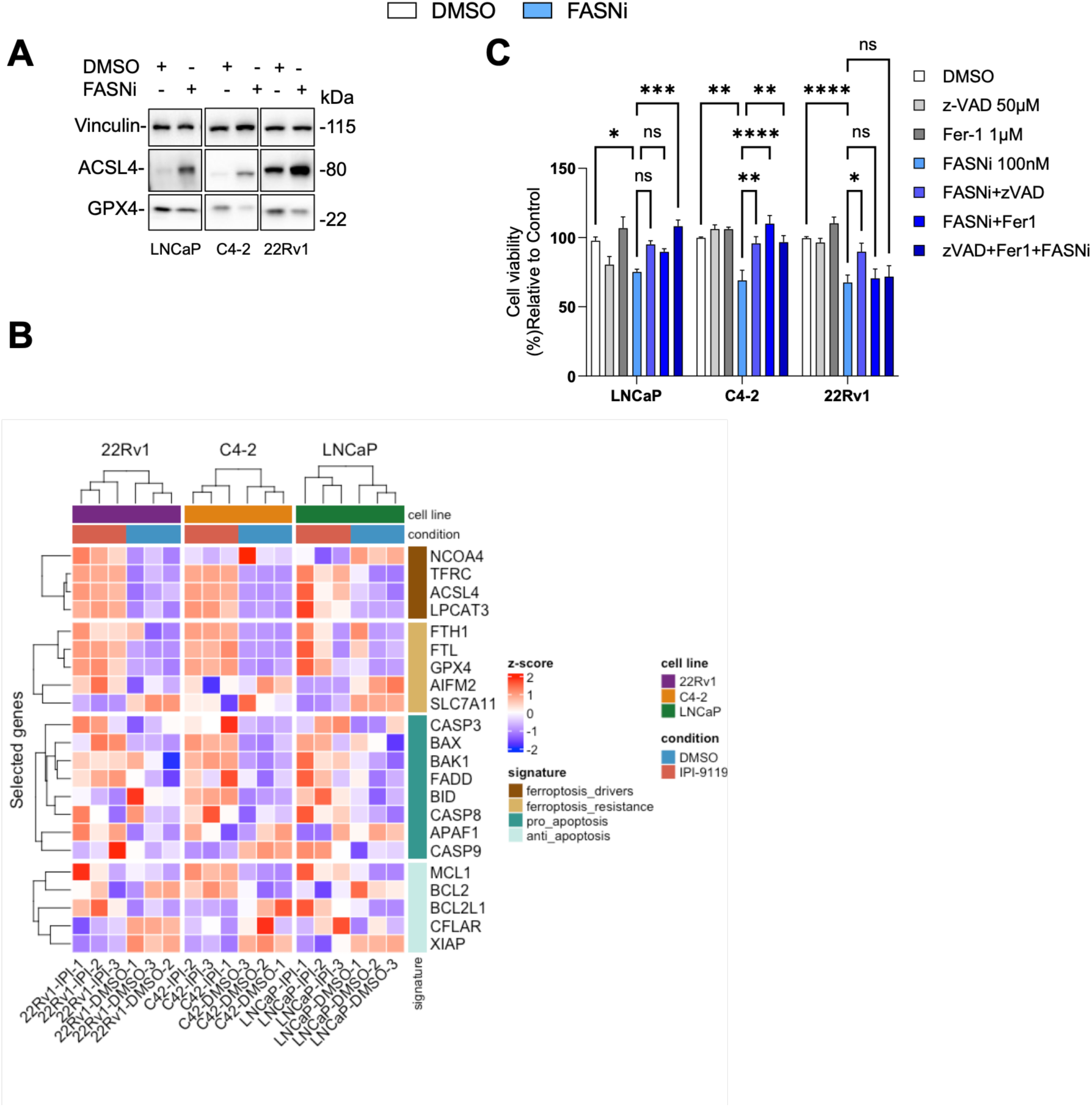
FASN inhibition sensitizes prostate cancer cells to lipid peroxidation. (A) Immunoblot analysis of ACSL4, and GPX4 following FASN inhibition. Vinculin served as a loading control. (B) Heatmap of regulated cell death–associated genes following IPI-9119 treatment compared with DMSO. (C) Cell viability measured using the CellTiter-Glo ATP-based luminescence assay following treatment with FASN inhibitor (IPI-9119, 100nM) alone or in combination with Ferrostatin-1 (10 μM), and/or Z-VAD-FMK (50μM), expressed as percentage relative to DMSO-treated control. Data represent mean ± SEM from three independent biological replicates (n = 3). Statistical analysis was performed using one-way ANOVA with Tukey’s multiple-comparison test. ns, not significant; **P < 0.01; ***P < 0.001; ****P < 0.0001.

### 4.3 Reduced SFA and increased PUFA foster oxidative damage

In addition to membrane phospholipids, FASN inhibition promoted the accumulation of PUFA in neutral lipid pools, with cholesteryl esters (CEs) acting as a primary reservoir for highly unsaturated fatty acids. CE formation is a key adaptive pathway for sequestering excess exogenous PUFAs while limiting their incorporation into peroxidation-prone membranes. Docosahexaenoic acid (DHA), an omega-3 PUFA containing 22 carbons and six double bonds (22:6) (Figure 4A), accumulated predominantly within CE pool, in both HS and CR cells. DHA was therefore used as the selected tracer. Uptake of labeled DHA was significantly increased in FASNi-treated cells compared to controls (Figure 4B). Importantly, however, when exogenous palmitate was simultaneously available, cells preferentially incorporated palmitate over DHA following FASN inhibition (Figure 4C).

**Figure 4.**
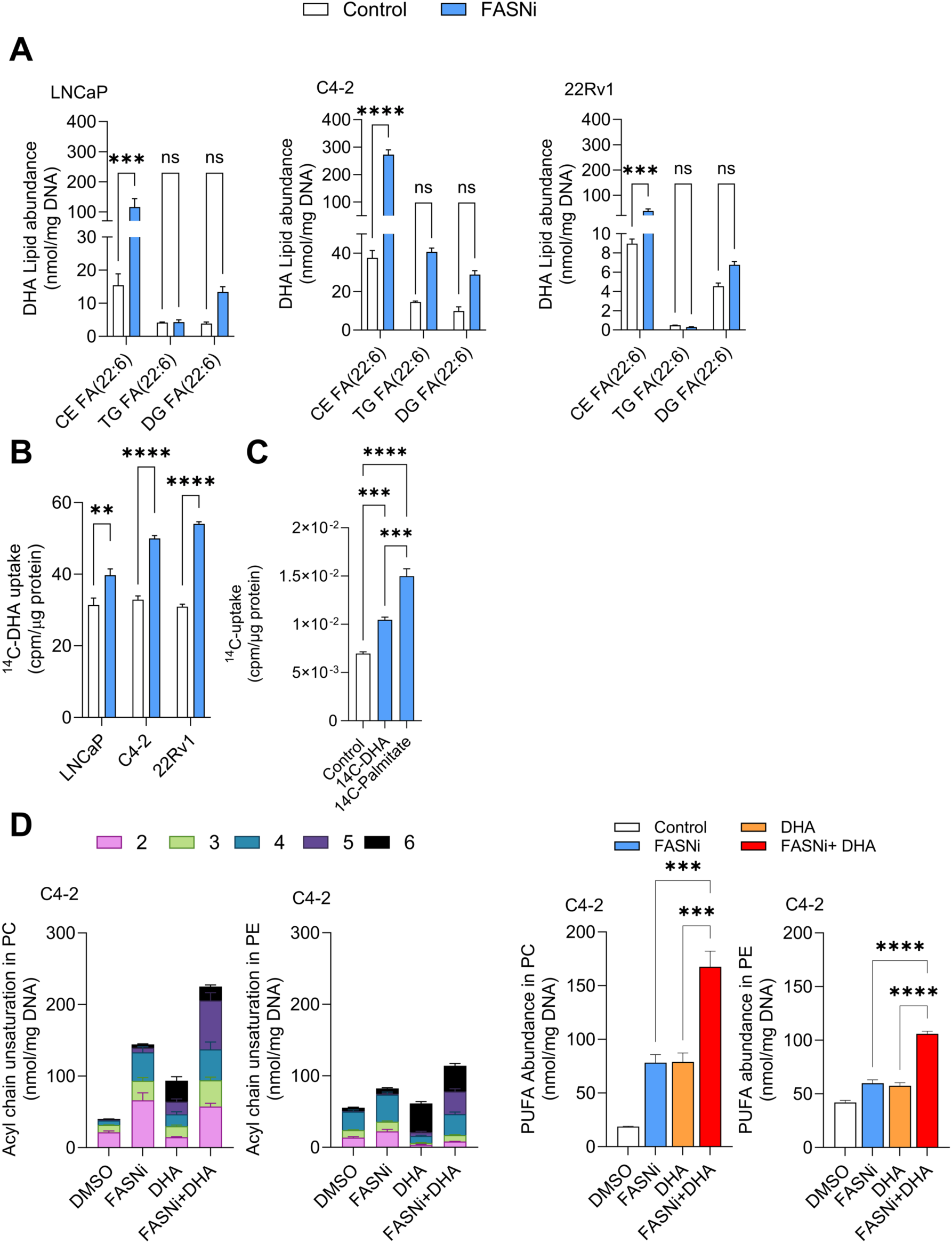
FASN inhibition increases exogenous DHA uptake and promotes PUFA-enriched phospholipid remodeling. (A) Abundance of DHA-containing lipid species across major lipid classes in LNCaP and C4-2 cells following FASN inhibition (IPI-9119 100 nM) for 6 days of treatment, expressed relative to control as indicated. (B) Uptake of [^14^C]-DHA in LNCaP and C4-2 cells following FASNi treatment, expressed as counts per minute (cpm) normalized to protein (µg). (C) Uptake of radiolabeled DHA or palmitate in C4-2 cells, expressed as cpm normalized to protein. (D) Phospholipid acyl-chain unsaturation distributions in C4-2 cells (PC and PE; stacked by number of double bonds) and quantification of PUFA-containing PC and PE species (≥2 double bonds) following treatment with DMSO, FASNi, DHA, or FASNi + DHA. Data represent mean ± SEM from three independent biological replicates (n = 3). Statistical analysis was performed using one-way ANOVA with Tukey’s multiple-comparison test. ns, not significant; *P < 0.05; **P < 0.01; ***P < 0.001; ****P < 0.0001.

We previously demonstrated that either FASNi alone^4,14^ or DHA treatment as a single agent^32,39^ increases oxidative damage in prostate cancer cells. Here, we investigated whether combining FASN inhibition with exogenous PUFA supplementation further enhances susceptibility to oxidative damage and cell death. Because FASNi treatment promotes PUFA incorporation into membrane phospholipids (Figure 2), we first analyzed phospholipid acyl chain unsaturation in PC and PE species under these conditions. Both FASNi and DHA, as single agents, increased the degree of unsaturation within membrane lipids. However, the combination treatment produced an even more pronounced increase in membrane unsaturation compared with either single agent (Figure 4D). This observation aligns with the elevated abundance of PUFA in PC and PE species (Figure 2A), which are associated with increased membrane oxidative sensitivity ^22,24,40^. We next assessed whether increased unsaturation sensitizes cells to mitochondrial stress. Intracellular superoxide levels increased following combined FASNi and DHA treatment relative to control and single-agent conditions (Figure 5A). In parallel, mitochondrial membrane potential analysis revealed marked hyperpolarization in the combination group compared with either single treatment, a phenotype previously associated with early mitochondrial redox stress during ferroptotic signaling (Figure 5A). Notably, palmitate supplementation reversed the hyperpolarization induced by the FASN inhibition treatment (Supplementary Figure 1A).

**Figure 5.**
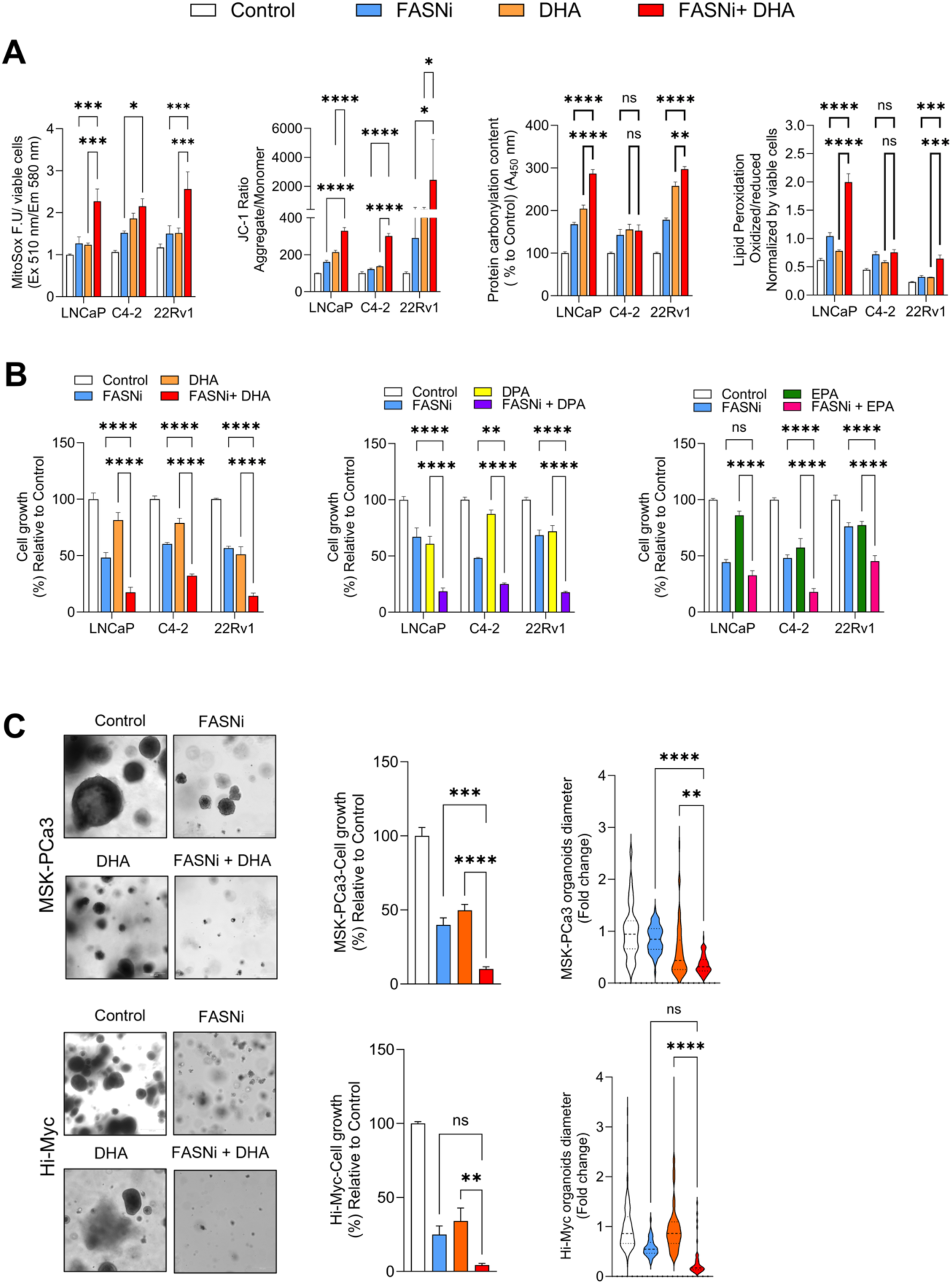
Exogenous PUFA potentiates the effects of FASN inhibition on oxidative stress and prostate cancer cell growth. (A) Oxidative stress markers in LNCaP, C4-2 and 22Rv1 cells treated with FASN inhibitor (IPI-9119, 100 nM), DHA (100 μM), or the combination for 6 days. Mitochondrial superoxide levels were measured using MitoSOX and normalized to viable cell number. Mitochondrial membrane potential was assessed using the JC-1 aggregate/monomer fluorescence ratio. Protein carbonylation is expressed as percentage relative to control. Lipid peroxidation was quantified using BODIPY 581/591 C11 and expressed as oxidized/reduced fluorescence ratio normalized to viable cell number. (B) Cell growth was assessed by direct cell counting using the Vi-CELL BLU automated cell counter following 6 days of treatment with FASNi (IPI-9119, 100nM), PUFAs (100 μM DHA; 100 μM DPA; 120 μM EPA), and their combinations. Growth is expressed as percentage relative to DMSO-treated control (C) Representative brightfield images of MSK-PCa3 and Hi-Myc organoids treated with FASNi, DHA, or the combination for 15 days, scale bar 300 μm. Quantification of organoid growth is expressed as a percentage relative to the control. Violin plots show organoid diameter fold change relative to control. Data represent mean ± SEM from three independent biological replicates unless otherwise indicated. Statistical analysis was performed using one-way ANOVA with Tukey’s multiple-comparison test. For clarity, statistical annotations indicate comparisons between the combination treatment and single-agent conditions. ns, not significant; **P < 0.01; ***P < 0.001; ****P < 0.0001.

To directly assess oxidative damage, we quantified protein carbonylation and lipid peroxidation (Figure 5A). Combined FASN inhibition and DHA treatment significantly increased oxidative damage markers in HS LNCaP and CR 22Rv1 cells, whereas CR C4-2 exhibited a more attenuated response, with no significant increase in lipid peroxidation under combination treatment. This differential oxidative stress response paralleled the more pronounced growth suppression observed in LNCaP cells (Figure 5B).

Considering the effects observed with the combination of FASN inhibition and DHA, particularly the elevation of intracellular ROS and increased lipid peroxidation, we next investigated whether structurally related PUFAs could elicit similar effects on cell proliferation (Figure 5B, left panel). In addition to DHA (22:6), we tested docosapentaenoic acid (DPA; 22:5) (Figure 5B, middle panel) and eicosapentaenoic acid (EPA; 20:5) (Figure 5C, right panel), two omega-3 long-chain PUFAs characterized by multiple unsaturation sites that confer susceptibility to peroxidation (Figure 5B). When administered as single agents, all three PUFAs reduced prostate cancer cell growth. Their combination with FASN inhibition produced a greater inhibitory effect than either single treatment, prompting validation in 3D organoid models. Consistent with the 2D findings, combined FASN inhibition and DHA significantly reduced growth in MSK-PCa3 and Hi-Myc organoids (Figure 5C). Together, these data support a model in which modulation of lipid composition enhances tumor vulnerability to oxidative stress, ultimately promoting cell death.

### 4.4 PUFA-enriched diet and FASN inhibition remodel systemic and tumoral lipidomes in Hi-Myc mice

MYC-driven prostate tumors exhibit increased dependency on DNL^41,42^, and incorporation of exogenous PUFAs enhances the cytotoxic effects of FASN inhibition *in vitro*. To determine whether a PUFA-enriched diet could amplify the therapeutic response to FASN blockade *in vivo*, we used the Hi-Myc genetically engineered mouse model (GEMM)^43,44^. After weaning (21 days), mice were randomized to receive either a High PUFA Diet (HPD; Teklad TD.230355) or an SFA/MUFA-enriched control isocaloric diet (HSD; Teklad TD.230313) (Table 1). Each dietary group was further stratified to receive FASNi (TVB-3664). FASNi was initiated at 7 weeks of age for the 12-week endpoint cohort and at 14 weeks of age for the 20-week endpoint cohort (Figure 6A). Body weights were comparable across all groups (Supplementary Figure S2A).

**Figure 6.**
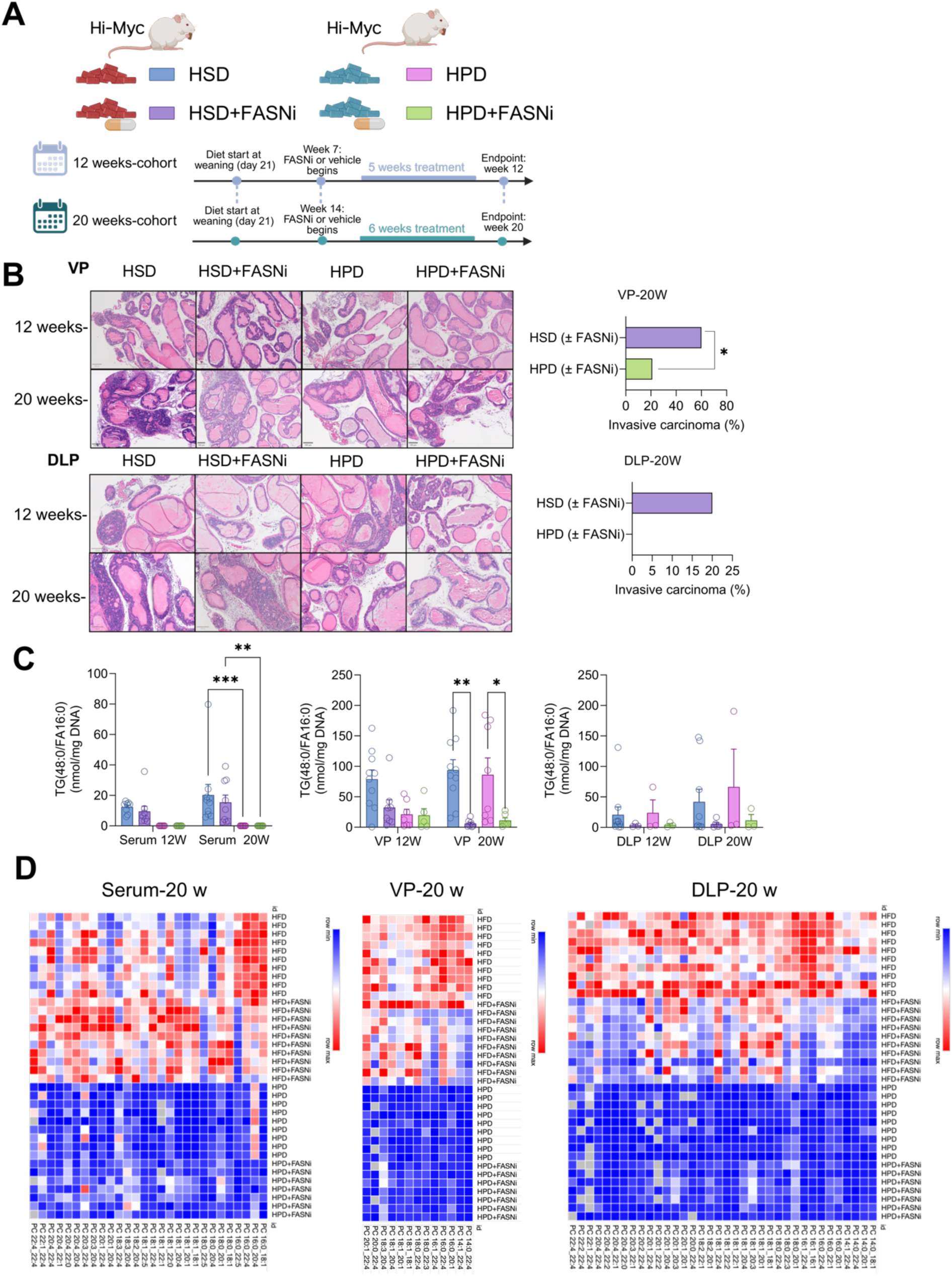
Dietary PUFA enrichment remodels systemic and tumor lipidomes and reduces invasive progression in Hi-Myc mice. (A) Animal model experimental design. (B) Representative images of the incidence of invasive adenocarcinoma in ventral (VP) and dorsolateral (DLP) prostate lobes of Hi-Myc mice fed a high saturated fat diet (HSD) or a PUFA-enriched diet (HPD) with or without FASN inhibitor (TVB-3664). Statistical comparisons were performed using Fisher’s exact test. (C) Tripalmitin (TG 16:0/16:0/16:0) abundance in serum, VP and DLP. (D) Heatmap of phosphatidylcholine (PC) modulation at 20 weeks in serum, VP, and DLP. Tissue lipid species were normalized to DNA content. Serum lipid species were normalized to volume. Data represent mean ± SEM (n = 8–10 mice per group as indicated). Statistical analysis for lipidomic comparisons was performed using Mann–Whitney U test or one-way ANOVA with appropriate multiple-comparison correction as described in Methods.

**Table 1.** Dietary composition. Table 1. Composition of murine diets. Purified, isocaloric diets were used throughout the study. Diets were matched for total caloric density and macronutrient distribution and differed primarily in fatty-acid composition. The high saturated and monounsaturated fatty acid diet (HFD) is enriched in saturated fatty acids (SFAs) and monounsaturated fatty acids (MUFAs), whereas the high polyunsaturated fatty acid diet (HPD) contains increased polyunsaturated fatty acids (PUFAs) derived from safflower oil and fish oil. Detailed fatty-acid distribution and percentage of total energy contribution are provided.

|  | HSD | HPD |
| --- | --- | --- |
| **Diet ID (Teklad)** | TD.230313 | TD.230355 |
| **Diet designation** | High SFA + MUFA (PO, O) | 55% PUFA (SO, FO, G) |
| **Protein (% kcal)** | 15 | 15 |
| **Carbohydrate (% kcal)** | 55 | 55 |
| **Fat (% kcal)** | 30 | 30 |
| **Energy density (kcal g<sup>-1</sup>)** | 4.1 | 4.1 |
| **Primary fat sources** | Palm oil | Safflower oil + fish oil |
| **Fatty-acid composition** | ~90% SFA + MUFA~10% PUFA | ~55% PUFA |
| **Protein source** | Casein | Casein |
| **Carbohydrate sources** | Corn starch, maltodextrin, sucrose | rch, maltodextrin, sucrose |
| **Antioxidant** | TBHQ (0.03 g kg<sup>-1</sup>) | TBHQ (0.03 g kg<sup>-1</sup>) |
| **Diet form** | Pellet or powder | Pellet or powder |
| **Storage conditions** | $\leq 4^{\circ}\text{C}$ | $\leq 4^{\circ}\text{C}$ |
| **Diet replacement** | ≥ twice per week | ≥ twice per week |
| **Color code** | Orange | Green |

As expected, Hi-Myc mice developed prostatic intraepithelial neoplasia (PIN) in both ventral (VP) and dorsolateral (DLP) lobes^43^. By 20 weeks, HSD-fed mice progressed to invasive adenocarcinoma in VP and DLP lobes at frequencies consistent with established tumor kinetics. In contrast, HPD-fed mice showed significantly reduced progression to invasive disease (21.4% (HPD+/-FASNi) vs 60% (HSD+/-FASNi); Fisher’s exact p = 0.038 in VP, while in DLP no invasive adenocarcinoma was found; Figure 6B), indicating that dietary PUFA enrichment alone restrains tumor progression. No invasive lesions were detected at 12 weeks in any group, and the anterior prostate remained free of invasive disease at both time points (Supplementary Figure S2B). Within the HSD background, FASN inhibition did not significantly reduce invasive incidence, since the high availability of dietary SFA/MUFAs likely compensated for DNL blockade. Similarly, under HPD, the protective effect of dietary PUFAs predominated, and additional FASN inhibition did not further decrease invasion, suggesting that histologic outcomes were predominantly diet-driven (Figure 6B).

To assess how dietary fat composition and FASN inhibition reshaped the metabolic landscape associated with these histologic outcomes, we performed lipidomic analyses of serum (Extended Data S2) and prostate tissues (Extended Data S3) from the ventral (VP) and dorsolateral (DLP) lobes at 12 and 20 weeks of age. Twelve-week serum, VP and DLP tissues results are presented in Supplementary Figures S3; the top three FAs altered at 20 weeks under each condition are summarized in Supplementary Table 1 and Extended Data S2-3.

A canonical DNL product, tripalmitin (TG 16:0/16:0/16:0)^25^, was markedly reduced and became undetectable in serum from HPD-fed mice, with or without FASNi. In prostate tissue, tripalmitin was significantly decreased in HSD + FASNi animals (Figure 6C), confirming effective *in vivo* DNL suppression. Dietary intervention induced broad lipidomic remodeling, particularly within serum phospholipids. HPD feeding was associated with globally reduced phosphatidylcholine (PC) (Figure 6D) and phosphatidylethanolamine (PE) abundance compared to HSD (Supplementary Figure S3), establishing dietary lipid composition as the dominant determinant of bulk membrane phospholipid pools.

FASNi induced depletion of PC and PE species, a signature that intensified at 20 weeks in HSD-fed animals but was attenuated under HPD, suggesting that exogenous PUFAs availability buffers the metabolic consequences of DNL blockade. Lipidomic changes in the DLP lobe paralleled those in VP but were less pronounced. Cross-compartment analysis at 20 weeks identified 72 lipid species consistently altered between HSD + FASNi and HPD + FASNi groups, including coordinated reductions in PC and PE species and reciprocal increases in lysophospholipids (LPC, LPE), with mixed alterations in PI, PS, and PG (Supplementary Table 1; Extended Data S2–S3). Notably, PC and PE abundance remained higher in HSD ± FASNi groups relative to HPD ± FASNi, reinforcing that dietary composition, rather than FASN inhibition alone, dictates bulk phospholipid abundance.

We next examined palmitate (C16:0) distribution within phospholipid acyl chains. In VP and DLP tissues, palmitate-containing phospholipid species were significantly reduced by FASNi, particularly under HSD (Supplementary Figure S4), consistent with suppression of endogenous palmitate synthesis. In contrast, serum phospholipid-associated palmitate remained largely unchanged across conditions. This divergence suggests that circulating phospholipid pools are predominantly shaped by dietary fatty acid influx and systemic remodeling processes, whereas intratumoral phospholipid palmitate content more directly reflects local DNL activity.

In serum, a significant reduction in total palmitate (FA 16:0), rather than phospholipid-bound palmitate, was observed specifically in the HSD + FASNi group (Figure Supplementary S4), underscoring compartment-specific metabolic regulation.

Given that palmitate is a substrate for elongation and desaturation, we quantified downstream acyl species. Lipids containing C16:1 were consistently reduced by FASNi, reaching statistical significance in HSD + FASNi versus HSD alone (Figure Supplementary S4). Moreover, C18:1-containing lipids were downregulated in both serum and prostate tissue (Figure Supplementary S4), indicating that inhibition of palmitate synthesis affects elongation/desaturation pathways.

## 6. Discussion

Lipid metabolism is profoundly reprogrammed in prostate cancer, and elevated DNL provides prostate cancer cells with FAs essential for cell membrane synthesis, mitochondrial function, and robust AR signaling^33,45^. FASN is frequently overexpressed in high-grade and castration-resistant tumors and is associated with poor prognosis and disease progression^8,46^. FASN activity fuels tumor growth^47^ and sustains AR signaling^4^, its pharmacological inhibition suppresses AR signaling and is currently under clinical evaluation with enzalutamide in metastatic CRPC (NCT05743621).

In this study, we demonstrate that pharmacological inhibition of DNL, combined with exogenous PUFA supplementation, suppresses prostate cancer growth *in vitro* and *in vivo.* This inhibition of growth has multiple underlying causes. Simultaneous restriction of endogenous SFA/MUFA supply increases incorporation of oxidation-prone PUFAs into membrane lipids. FASN inhibition also disrupts mitochondrial ATP-linked respiration in HS and CR prostate cancer cells, thereby reducing mitochondrial oxidative capacity. Consistent with our findings, previous studies using TVB-2640, cerulenin, and orlistat have shown that pharmacological FASN blockade impairs mitochondrial respiration, including NADH- and succinate-linked substrate oxidation, thereby reducing OXPHOS capacity and ATP production^48,49^. Finally, from an energetic standpoint, DNL blockade primarily impairs oxidative metabolism without a compensatory increase in ECAR, and therefore no glycolytic shift.

Rather than representing a generalized metabolic collapse, this bioenergetic phenotype likely reflects a selective vulnerability of AR-driven tumors to DNL inhibition. Although prostate tumors can maintain growth under lipid restriction by increasing glucose utilization^50^, the absence of a compensatory ECAR increase in our models indicates that FASN-inhibited cells fail to engage this adaptive response. By simultaneously depriving cells of endogenously synthesized SFA/MUFA and increasing PUFA incorporation, we generated a membrane lipid landscape with heightened susceptibility to peroxidation.

This shift destabilized redox homeostasis, manifested as mitochondrial hyperpolarization, ROS accumulation, and elevated lipid and protein oxidation, and was associated with growth suppression and activation of multiple cell-death pathways. We recently reported that FASN blockade induces DNA damage in prostate cancer cells, driven at least in part by accumulation of sphingomyelins and PUFA-enriched sphingolipids, which exacerbate lipid peroxidation-associated oxidative stress and compromise genomic integrity^14^.

Notably, DNL blockade has also been shown to impair ER stress responses in prostate cancer^4^ and multiple myeloma cells^35^, while DHA itself can induce ROS production and oxidative damage^32,39,51^. In addition, FASN inhibition increased ACSL4 expression, which promotes PUFA incorporation into phospholipids and is associated with ferroptosis sensitivity. ACSL4 induction, together with partial rescue by ferrostatin-1, supports a contribution of lipid peroxidation-dependent cell death. This mechanistic complexity may have therapeutic implications, as it suggests potential benefit from combining FASN inhibitors with agents that modulate ferroptosis sensitivity, apoptotic priming, or ER-stress.

A diet enriched in PUFAs and lower in saturated fats, significantly reduced invasive carcinoma burden in an autochthonous GEMM, Hi-Myc, previously shown to be DNL-dependent^42,52^. Importantly, FASN inhibition did not further reduce invasive incidence within the HPD background, suggesting that the dietary effects predominated over pharmacologic intervention *in vivo*. This lack of additional antitumor effect is unlikely to reflect an absence of FASN inhibitor activity, as supported by reduced tripalmitin levels. PUFA enrichment, particularly with long-chain omega-3 fatty acids such as DHA, partially phenocopies FASN inhibition by directly suppressing the endogenous lipogenic program. Consistent with this, we previously showed that DHA downregulates enzymes involved in fatty acid synthesis, desaturation and elongation, decreases FASN protein expression, reduces *de novo* lipid synthesis, and suppresses AR-FL and AR-V7 in prostate cancer models^32^. Thus, in a PUFA-enriched dietary context, endogenous lipogenesis and AR-dependent growth programs may already be attenuated. Conversely, the lack of a significant effect of FASN inhibition within the HSD background may reflect metabolic compensation by exogenous dietary fatty acids, in that abundant circulating SFA/MUFA substrates bypass the requirement for tumor-intrinsic DNL and preserve lipid-dependent processes required for tumor progression. In keeping with these findings, high-fat diets, particularly enriched in SFA/MUFAs, are linked to increased aggressiveness and poorer outcomes for prostate cancer^15,41,53–56^, whereas diets enriched in long-chain PUFAs, particularly omega-3, have been associated with antitumor effects^32,57^, at least in part by altering membrane lipid composition and downstream signaling pathways and by accelerating AR degradation^58^.

Lipidomic profiling revealed a convergence between pharmacologic FASN blockade and dietary intervention, with consistent depletion of phospholipids enriched in MUFA chains, particularly those derived from *de novo* palmitate synthesis (e.g., PC 16:1_16:1, PC 16:1_20:3). These data reinforce that MYC-driven prostate tumors maintain a dependency on endogenously synthesized palmitate not merely as a bulk lipid source, but to sustain a defined subset of phospholipid species critical for membrane organization and signaling. Recent studies demonstrate that cancer cells require SFA/MUFAs to maintain lipid raft function^59^ and to buffer oxidative stress^60^, supporting this interpretation. Together, these findings indicate that dietary lipid composition can modulate tumor lipid pools and, under defined metabolic contexts, potentially sensitize tumors to DNL blockade.

The mechanistic link between membrane lipid remodeling, redox imbalance, and cell-death programming highlights several opportunities for clinical translation. First, lipidomic signatures, including tripalmitin depletion and changes in palmitate-derived SFA/MUFA species, provide candidate pharmacodynamic biomarkers of FASN target engagement, whereas circulating lipid profiles report dietary exposure. Second, the enhanced growth suppression observed with combined FASN blockade and PUFA supplementation indicates that the response to FASN-targeted therapy is conditioned by extracellular fatty acid composition. Together, these measurable biomarkers provide a framework for integrating precision nutrition with pharmacologic intervention.

Our findings support three central conclusions: First, the therapeutic consequences of FASN inhibition are conditioned by extracellular fatty acid availability: PUFAs expose the oxidative vulnerability created by DNL blockade, whereas SFA/MUFA availability can support metabolic compensation. Second, this vulnerability is executed through overlapping lipid peroxide-dependent and caspase-dependent death programs, implicating both ferroptosis and apoptosis. Third, circulating and tumor lipidomic signatures of dietary exposure and FASN target engagement provide a measurable framework for incorporating precision nutrition into FASN-directed therapy.

## 7. Conclusion

Together, our findings support a metabolic model in which DNL blockade increases tumor dependence on exogenous fatty acids, creating vulnerability to oxidative stress that is determined by the composition of the extracellular lipid pool. Dietary lipid composition can substantially influence tumor progression and modulate the response to metabolic therapy. In vivo, FASN inhibition created a metabolic context in which SFA/MUFA availability supported metabolic compensation, whereas PUFA enrichment increased tumor vulnerability. Specifically, FASN blockade lowers palmitate and tripalmitin and suppresses SFA/MUFA-supported AR and mitochondrial functions, while concomitant PUFA enrichment enriches membranes with highly oxidizable acyl chains, enhancing oxidative stress and reducing tumor cell growth through overlapping ferroptotic and apoptotic mechanisms. In vivo, dietary PUFA enrichment reshapes systemic and prostate lipidomes and attenuates disease progression in MYC-driven prostate cancer. Notably, this dietary intervention can equal or exceed the impact of pharmacological FASN inhibition, emphasizing that nutrient availability is an active determinant of therapeutic response to metabolic therapies. By exploiting lineage-specific lipid dependencies, the combination of DNL inhibition and dietary modulation may define a therapeutically actionable metabolic window in prostate cancer. Finally, circulating and tumor lipidomic signatures, including tripalmitin depletion and changes in palmitate-derived lipid species, provide candidate biomarkers of dietary exposure and FASN target engagement, supporting a precision nutrition framework for patient stratification and treatment monitoring.

## 8. Acknowledgements

This work and the contributing investigators were supported by Prostate Cancer Foundation (PCF-22CHAL05, PCF 22CHAL13, and 25YOUN19), Department of Defense (DoD-W81XWH-19-1-0566, HT9425-25-1-0385), National Institute of Health (P01CA265768, P50CA211024, R01CA234162), American Cancer Society (PF-24-1318851-01-MM), American Italian Cancer Foundation Postdoctoral Fellowships, Associazione Italiana per la Ricerca sul Cancro (AIRC) “Ezio, Maria e Bianca Panciera” Fellowship for Abroad, Research Foundation Flanders (FWO) grant G032525N, Belgian Foundation Against Cancer (Stichting tegen Kanker) F/2020/1417, Kom Op Tegen Kanker (Stand Up to Cancer), Leuven Future Fund LISCO-BIOMED. We are grateful to Weill Cornell Medicine vivarium staff for their great support in animal studies. Also, we thank Dr. Yu Chen (Memorial Sloan-Kettering Cancer Center, New York, NY, USA), who provided the MSK-PCa3 organoids, and Lipometrix (Leuven, Belgium) for the lipidomic data. We also acknowledge Robert Willumstad for his support.

## 9. Conflict of Interest

The authors declare no conflict of interest

## 10. Declaration of generative AI and AI-assisted technologies in the writing process

During the preparation of this work, the author(s) used ChatGPT in the writing process in order to only improve the readability and language of the manuscript. After using this tool/service, the author(s) reviewed and edited the content as needed and take(s) full responsibility for the content of the published article.

## Notes

### Competing Interest Statement

The authors have declared no competing interest.

### Summary of Updates

This version has been revised to incorporate additional experimental data and improve the organization and clarity of the manuscript. The castration-resistant prostate cancer cell line 22Rv1 has now been included in the main figures, and the figure order has been reorganized accordingly. Two new datasets have also been added to the revised Figure 3: a heatmap highlighting regulated cell death-associated genes altered following FASN inhibition and a cell viability rescue experiment. The in vivo section has been expanded to include results from both the 12- and 20-week time points, including serum analyses and measurements of the ventral and dorsolateral prostate lobes. In addition, part of the lipidomic dataset has been transferred to the Supplementary Figures to improve the clarity and readability of the main figures. A new co-author has also been added to the revised version to reflect their contribution to the expanded work.

