## Supplementary Figures and Legends for "Suppression of de novo lipogenesis and dietary PUFA supplementation inhibit prostate cancer progression"

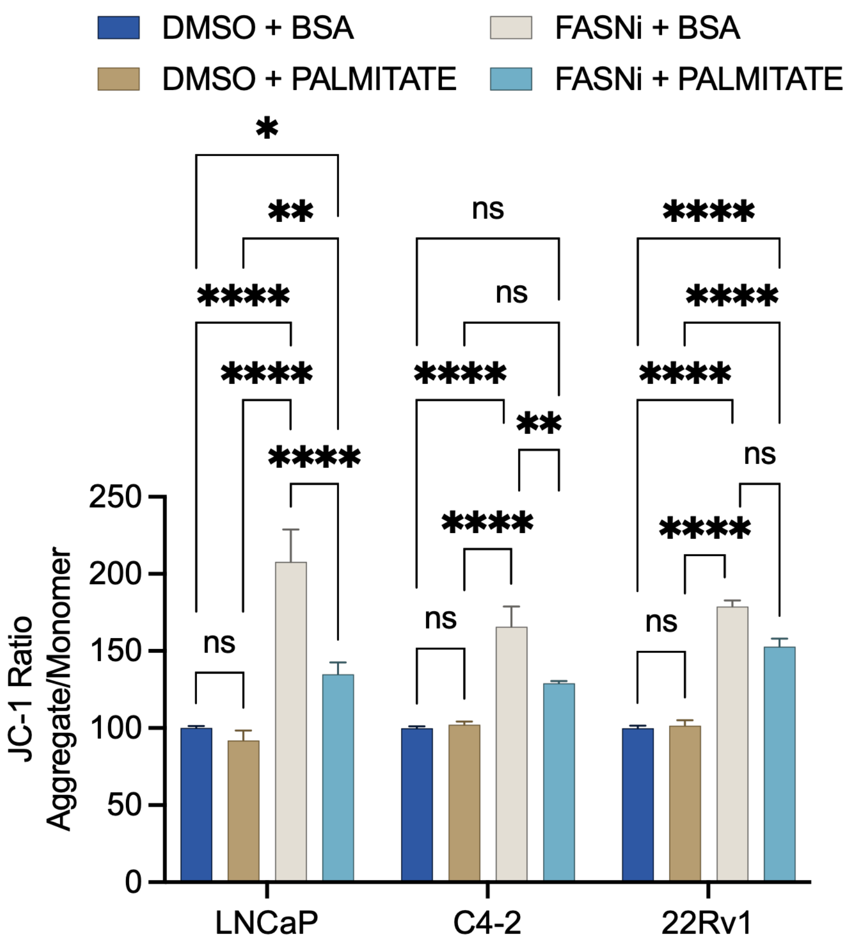


**S1. Exogenous palmitate reverses FASN inhibition-induced mitochondrial membrane hyperpolarization.** JC-1 fluorescence assay measuring mitochondrial potential in LNCaP, C4-2, and 22Rv1 cells treated with vehicle, palmitate, FASNi (IPI-9119 100 nM), or FASNi + palmitate. Cells were incubated with treatments for 6 days. Data represent mean ± SEM from n = 3 independent biological experiments. Statistical analyses were performed using two-way ANOVA with Tukey or Šidák post hoc comparisons; significant differences are indicated as *$P < 0.05$, **$P < 0.01$, ****$P < 0.0001$.

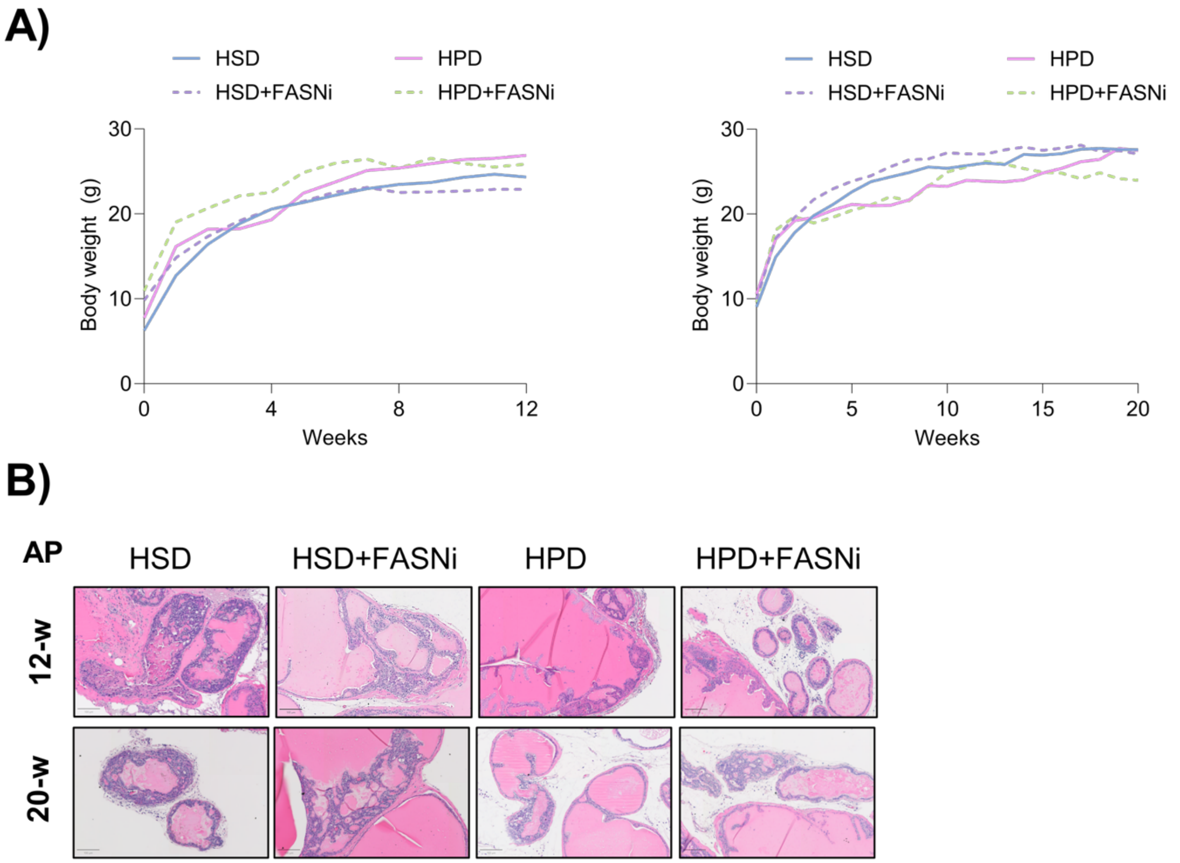


**S2. Body weight curves and histopathology of anterior prostate lobes in Hi-Myc mice under dietary and FASN inhibition intervention.** (A) Body weight over time of Hi-Myc mice fed a high-saturated and monounsaturated diet (HSD), or a high-polyunsaturated diet (HPD), each with or without FASN inhibitor (FASNi, TVB-3664). Left, 12-week cohort; right 20-week cohort. (B) Representative H&E-stained sections of the anterior prostate (AP) at 12 and 20 weeks for the HSD, HSD+FASNi, HPD, and HPD+FASNi groups.

S3

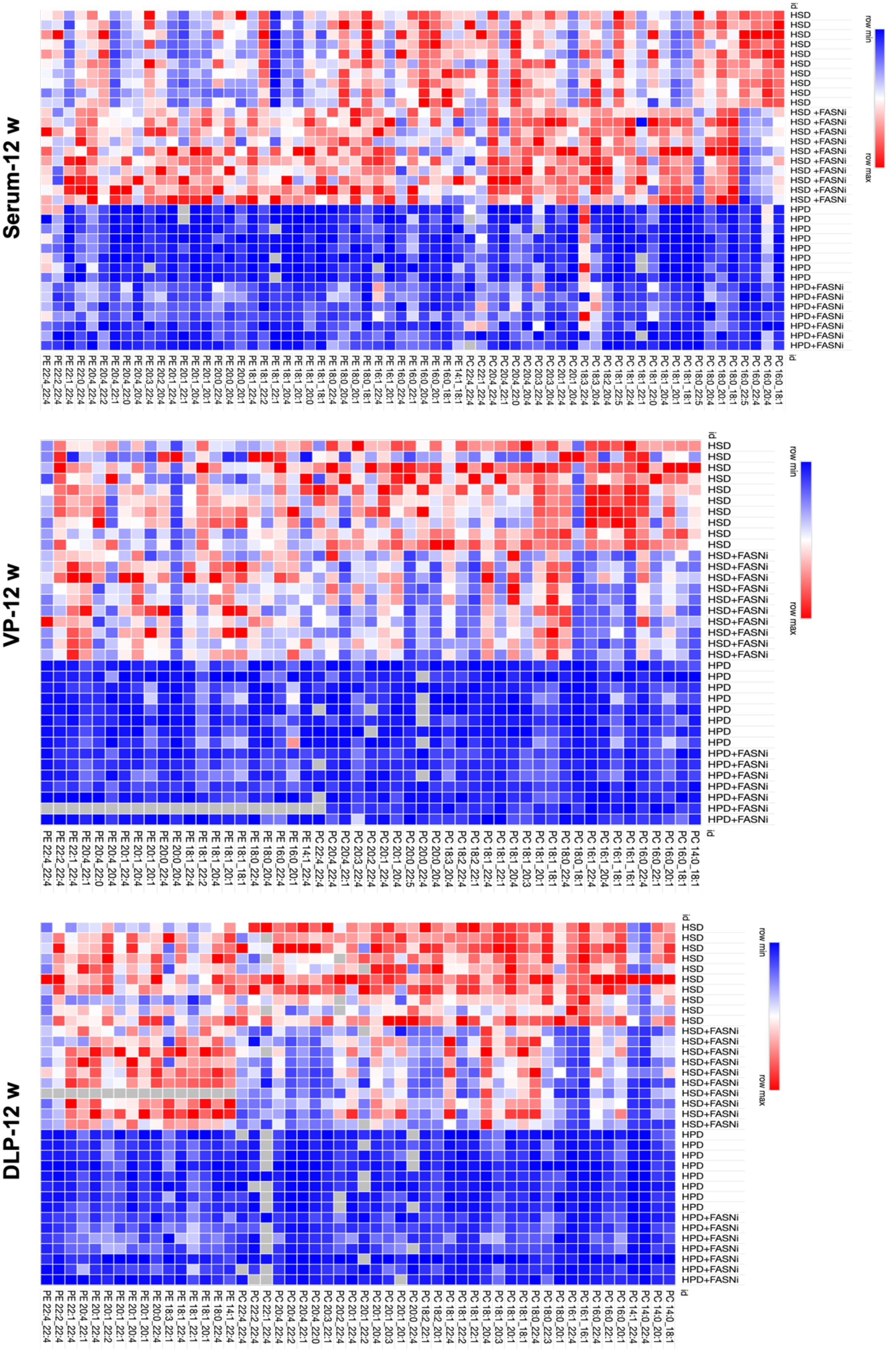

Serum-12 w
VP-12 w
DLP-12 w
row min
row max
HSD
HSD +FASNi
HSD+FASNi
HPD
HPD+FASNi

**S3. Global phospholipid remodeling in prostate tissue under dietary lipid modulation and FASN inhibition.** Heatmaps of phospholipid species abundance in serum and ventral prostate (VP) and dorsolateral prostate (DLP) lobes at 12 weeks**.** Columns represent lipid species, and rows represent individual biological samples. Color scale indicates row-normalized abundance. Heatmaps were generated using Morpheus, https://software.broadinstitute.org/morpheus

*Table 1. Differential lipid species abundance and leading fatty-acid alterations across compartments at 20 weeks*

| Comparison (20 weeks) | Compartment | ↓ Species (n) | ↑ Species (n) | Top 3 main regulated FA* |
|---|---|---|---|---|
| HSD vs HSD + FASNi | Serum | 134 | 119 | ↓ 16:0, 18:0, 22:5 ; ↑18:1, 20:4, 20:1 |
| | VP | 526 | 317 | ↓ 18:1, 16:0, 20:4 ; ↑ 20:5, 18:1, 18:2 |
| | DLP | 131 | 7 | ↓ 16:0, 18:0, 18:2 ; ↑ 18:1, 20:1, 20:2 |
| HPD vs HPD + FASNi | Serum | 62 | 117 | ↓ 16:0, 22:6, 18:0 ; ↑ 18:1, 18:2, 16:1 |
| | VP / DLP | – | – | – |
| HSD vs HPD | Serum | 412 | 556 | ↓ 18:1, 16:0, 20:3 ; ↑ 22:6, 20:5, 18:2 |
| | VP | 214 | 121 | ↓ 22:4, 20:4, 18:1 ; ↑ 20:5, 18:2, 22:6 |
| | DLP | 198 | 113 | ↓ 22:4, 20:4, 16:0 ; ↑ 20:5, 18:0, 18:2 |
| HSD + FASNi vs HPD + FASNi | Serum | 254 | 499 | ↓ 18:1, 20:4, 22:4 ; ↑ 22:6, 18:2, 20:5 |
| | VP | 90 | 59 | ↓ 22:4, 18:1, 20:1 ; ↑ 22:6, 20:5, 16:0 |
| | DLP | 111 | 131 | ↓ 22:4, 20:1, 20:4 ; ↑ 20:5, 18:2, 18:0 |

**Table 1.** Summary of lipid species significantly altered across serum and prostate lobes in response to diet and FASN inhibition. Comprehensive summary of lipid species significantly increased or decreased in serum, ventral prostate (VP) and dorsolateral prostate (DLP) under high saturated and monounsaturated fatty acid diet (HSD) or high polyunsaturated fatty acid diet (HPD), with or without FASN inhibition. For each comparison, the three most significantly altered fatty-acid species (based on adjusted P value and magnitude of change) are listed. Numbers indicate lipid species significantly decreased (↓) or increased (↑) in each pairwise comparison at 20 weeks. Statistical significance was determined using an unpaired two-tailed t-test with Welch's correction (no assumption of equal variance). Multiple testing correction was applied using a two-stage step-up procedure (Benjamini, Krieger and Yekutieli) with a false discovery rate (FDR) threshold of Q = 1%.

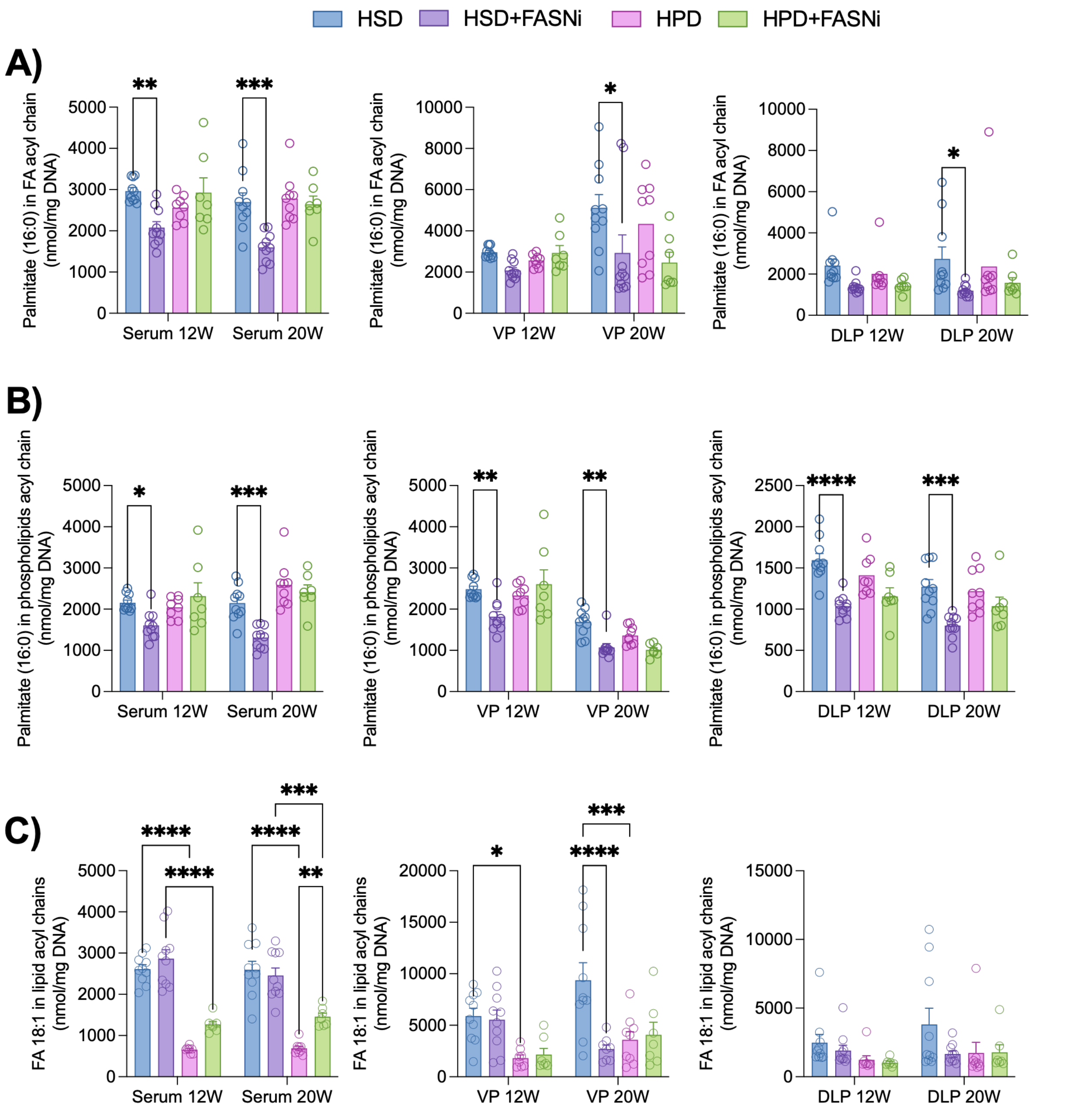


**S5. Fatty acid incorporation into lipid classes across diet and FASN inhibition conditions.** A) Palmitate levels into phospholipid acyl chains in ventral prostate, serum, and DLP. B) Palmitate levels into total fatty acid pools. C) Total FA (18:1) content in lipid acyl chains across tissues and time points. Fatty acid abundance was calculated from the summed acyl chains within annotated lipid species. Tissue values were normalized to DNA content; serum values to volume. Data represent mean ± SEM (n = 8–10 mice per group). Statistical significance was assessed using two-way ANOVA with Šidák correction. Each dot represents an individual mouse. Bars indicate mean ± SEM.

Statistical comparisons by two-way ANOVA with Tukey’s multiple comparison test. P*<0.05, P**<0.005, P***<0.0002, P****<0.0001
